# Predicting failures of Molteno and Baerveldt glaucoma drainage devices using machine learning models

**DOI:** 10.1101/646885

**Authors:** Paul Morrison, Maxwell Dixon, Arsham Sheybani, Bahareh Rahmani

**Affiliations:** Fontbonne University, Mathematics and Computer Science Department, St. Louis, MO, USA; Washington University, Department of Ophthalmology and Visual Sciences, St. Louis, MO, USA

## Abstract

The purpose of this retrospective study is to measure machine learning models’ ability to predict glaucoma drainage device failure based on demographic information and preoperative measurements. The medical records of sixty-two patients were used. Potential predictors included the patient’s race, age, sex, preoperative intraocular pressure, preoperative visual acuity, number of intraocular pressure-lowering medications, and number and type of previous ophthalmic surgeries. Failure was defined as final intraocular pressure greater than 18 mm Hg, reduction in intraocular pressure less than 20% from baseline, or need for reoperation unrelated to normal implant maintenance. Five classifiers were compared: logistic regression, artificial neural network, random forest, decision tree, and support vector machine. Recursive feature elimination was used to shrink the number of predictors and grid search was used to choose hyperparameters. To prevent leakage, nested cross-validation was used throughout. Overall, the best classifier was logistic regression.

## 1. Introduction

Glaucoma drainage devices are typically utilized in the management of glaucoma refractory to maximal medical therapy or prior failed glaucoma surgery. The devices can be divided into two categories: non-valved (e.g. Molteno and Baerveldt) and valved (e.g. Ahmed). Non-valved GDDs have been shown to be more effective in lowering intra-ocular pressure (IOP) and have lower rates of reoperation than valved GDDs, but experience more frequent failure leading to dangerously low IOP or reduction of vision to the point of absolute blindness [1]. However, there have been no studies directly comparing the two main types of non-valved GDDs despite their significantly different device profiles and implantation technique.

The accuracy of machine learning models in predicting GDD outcomes based on a minimal feature set provides a unique strategy to understand differences between these devices. Previous studies have predicted individual outcomes for other ophthalmic surgeries using machine learning and linear regression. Achiron et al. used extreme gradient boosted decision forests to predict the efficacy (final Visual Acuity (VA) divided by starting VA) of refractive surgery [2]. Rohm et al. compared five algorithms to predict postoperative VA at 3 and 12 months in patients with neovascular age-related macular degeneration [3]. Valdes-Mas et al. compared an artificial neural network with a decision tree to predict the occurrence of astigmatism and found the neural network superior [4]. Mohammadi et al. used neural networks to predict the occurrence of posterior capsule opacification after phacoemulsification [5]. Gupta et al. used linear regression to determine post-operative visual acuity based on patient demographics and pre-operative predictors [6]. Koprowski et al. compared hundreds of artificial neural network topologies to predict corneal power after corneal refractive surgery [7]. McNabb et al. used OCT (optical coherence tomography) to predict corneal power change after laser refractive surgery [8]. When comparing the Molteno and Baerveldt GDDs, demographic predictors included race, sex, and age at surgery. A total of seven clinical predictors were considered including:

### Implant Type

Identified by type of implant (Molteno or Baerveldt) and implant plate surface area.

### VA (logMAR)

“Visual Acuity (Logarithm of the Minimum Angle of Resolution).” A more reproducible visual acuity measurement often used in research. As Snellen visual acuity is more often collected in the clinic setting, conversion to logMAR allows easier statistical analysis.

### IOP

Intraocular pressure. Elevated IOP is the major risk factor for development of glaucoma.

### Number of medications

These include usage of beta-blockers, alpha-adrenergic agonists, prostaglandin analogs, or carbonic anhydrase inhibitors (CAI). The number of medications was calculated from patient records at each visit.

### Number of previous surgeries

Glaucoma drainage implants are typically placed after less-invasive treatments fail but may incidentally be utilized following other ophthalmic surgeries (e.g. phacoemulsification of cataracts or retinal surgeries).

### Type of previous surgeries

These include phacoemulsification or extracapsular cataract extraction (ECCE), trabeculectomy, pars plana vitrectomy, penetrating keratoplasty, Ex-PRESS shunt, iStent, or diode laser cyclophotocoagulation (dCPC).

### Diagnosis

Causes for glaucoma included open-angle, neovascular, uveitic, angle-closure, secondary to trauma, secondary to Penetrating Keratoplasty (PKP), pseudoexfoliation, and combined mechanism.

## 2. Data Description

Of 62 patients analyzed, 26 (41%) were determined to have device failure. The mean (± SD) follow-up time was 573±245.45 days (range 133-1037 days). Patient samples were balanced between male and female. White race was three times more common than black race and there was only one Asian patient (Tables 1 and 2). Implant failure was defined as final IOP was greater than or equal to 18, less than 20% reduction from pre-operative levels, or if repeat surgery for glaucoma was required (this did not include in-clinic procedures that did not indicate failure of the device itself). By the last recorded appointment, 35% (22) of patients had a failing IOP and 19% (12) required additional surgery. No patients in this group experienced loss of light perception.

**Table 1.**
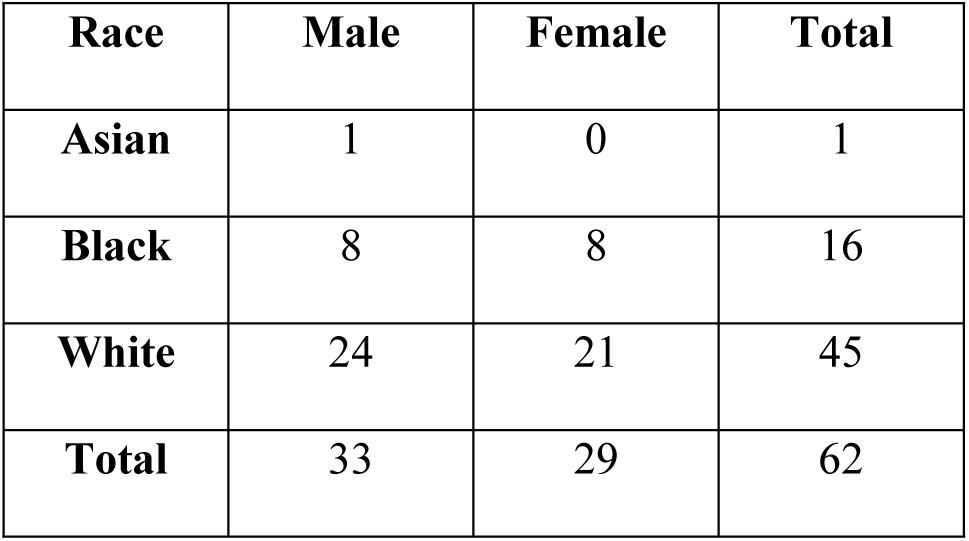
Number of participants by race and sex.

**Table 2.**
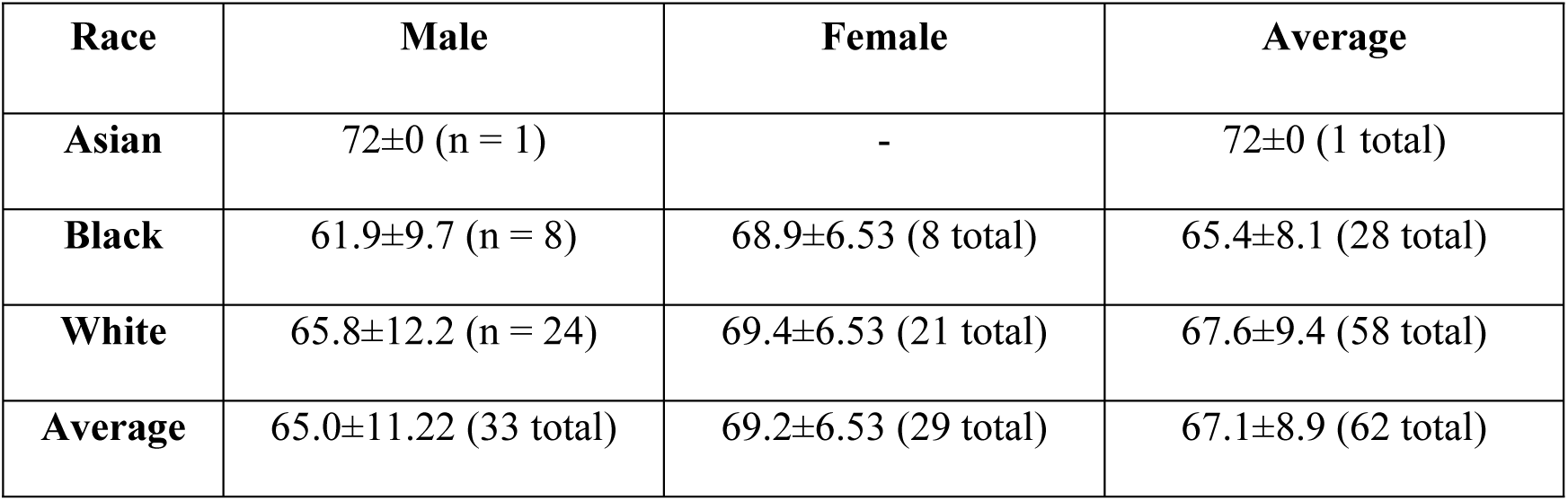
Average Age (mean ± SD) at Surgery by Race and Sex.

| Race | Male | Female | Average |
| --- | --- | --- | --- |
| Asian | 72±0 (n = 1) | - | 72±0 (1 total) |
| Black | 61.9±9.7 (n = 8) | 68.9±6.53 (8 total) | 65.4±8.1 (28 total) |
| White | 65.8±12.2 (n = 24) | 69.4±6.53 (21 total) | 67.6±9.4 (58 total) |
| Average | 65.0±11.22 (33 total) | 69.2±6.53 (29 total) | 67.1±8.9 (62 total) |

A total of 42 patients received a Baerveldt GDD (67%) and 20 received a Molteno GDD (27%). Forty-eight (77%) patients had surgery prior to placement of a GDD. Twelve patients (19%) required repeat surgery after initial placement of a GDD. Open-angle glaucoma was the most common underlying diagnosis (61%) with combined mechanism (11%) and chronic angle-closure (8%) being less common. There were also individual patients with either neovascular, uveitic, traumatic, or pseudoexfoliation glaucoma. A diagnosis of “Other” was given for 8% of the patients, which indicated a singular diagnosis was not able to be determined from chart review.

## 3. Methodology

All models in this study were validated using three-fold stratified cross validation, and all but the neural net were developed using recursive feature elimination and grid search. To prevent data leakage, final validation, grid searching, and feature selection were performed in separate cross validation loops, as recommended by Krstajic et al. [9]. In the outer loop, the final model was tested; in the middle loop, the best hyper parameters were chosen; and in the inner loop, the best feature subsets were selected. Within each loop, three-fold stratified cross-validation was used. Scaling and centering for continuous variables was performed as part of the model fitting procedure. The logistic regression, support vector machine, decision tree, and random forest classifiers were implemented in Python using Scikit-Learn [10], and the neural network classifier was implemented in R [11] using the caret package [12]. Accuracy was measured using classification accuracy and the area under the receiver operating characteristic curve (Area Under the Curve, or AUC).

### 3.1. Logistic Regression

Logistic regression, traditionally used for modeling, determines the class of each input observation by multiplying each feature by a constant, adding a bias term, and applying the logistic function. Any outcome above 0.5 is rounded to 1; any outcome below 0.5 is rounded to 0. The optimal logistic regression classifier used L2 regularization and a C parameter of 1, and had an accuracy score of 0.66±0.08 and a AUC value of 0.67±0.08. Based on the coefficients of the logistic regression model shown in Table 3 and Fig 1, Black race and initially taking beta-blockers are associated with implant failure.

**Table 3.** Feature Coefficients of Logistic Regression Classifier.

| Feature | Sign | Coefficient |
| --- | --- | --- |
| Black race | + | 0.83 |
| Initially taking Beta Blockers | + | 0.63 |
| White | - | 0.37 |
| Initially taking CAI | - | 0.36 |
| Baerveldt 350 mm<sup>2</sup> | - | 0.28 |

**Fig 1.**
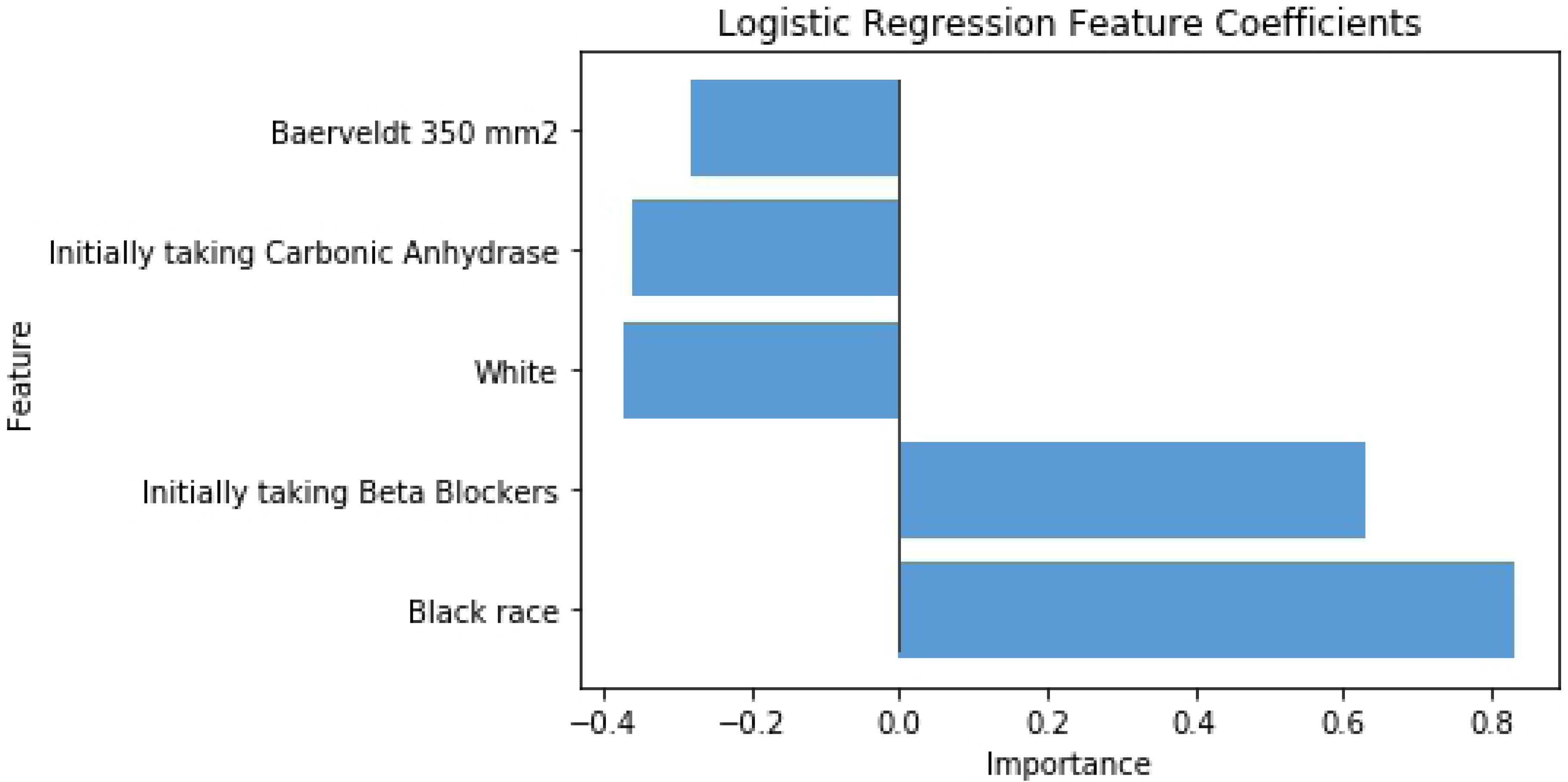
Feature Coefficients of Logistic Regression Classifier. The most important were black race and initially taking beta blockers.

### 3.2. Support Vector Machine (SVM)

A Support Vector Machine uses several data points (support vectors) to find the hyperplanes separating data classes that allow identification of a hyperplane giving the maximum margin [13]. The best classifier had a cost parameter of 0. Table 4 and Fig 2 show the feature coefficients of SVM classifier.

**Table 4.** Feature Coefficients of SVM Classifier.

| Feature | Sign | Coefficient |
| --- | --- | --- |
| Combined tube placement and phacoemulsification | - | 2 |
| Black race | + | 1.41 |
| No previous surgeries` | - | 1 |
| Baerveldt 350 mm<sup>2</sup> | - | 0.7 |
| Previous Phacoemulsification or ECCE | - | 0.6 |
| Initially taking beta blockers | + | 0.59 |
| Primary Open-Angle Glaucoma | + | 0.49 |
| Previous Trabeculectomy | - | 0.36 |
| Initially taking CAI | - | 0.34 |
| Initial VA (logMAR) | + | 0.28 |

**Fig 2.**
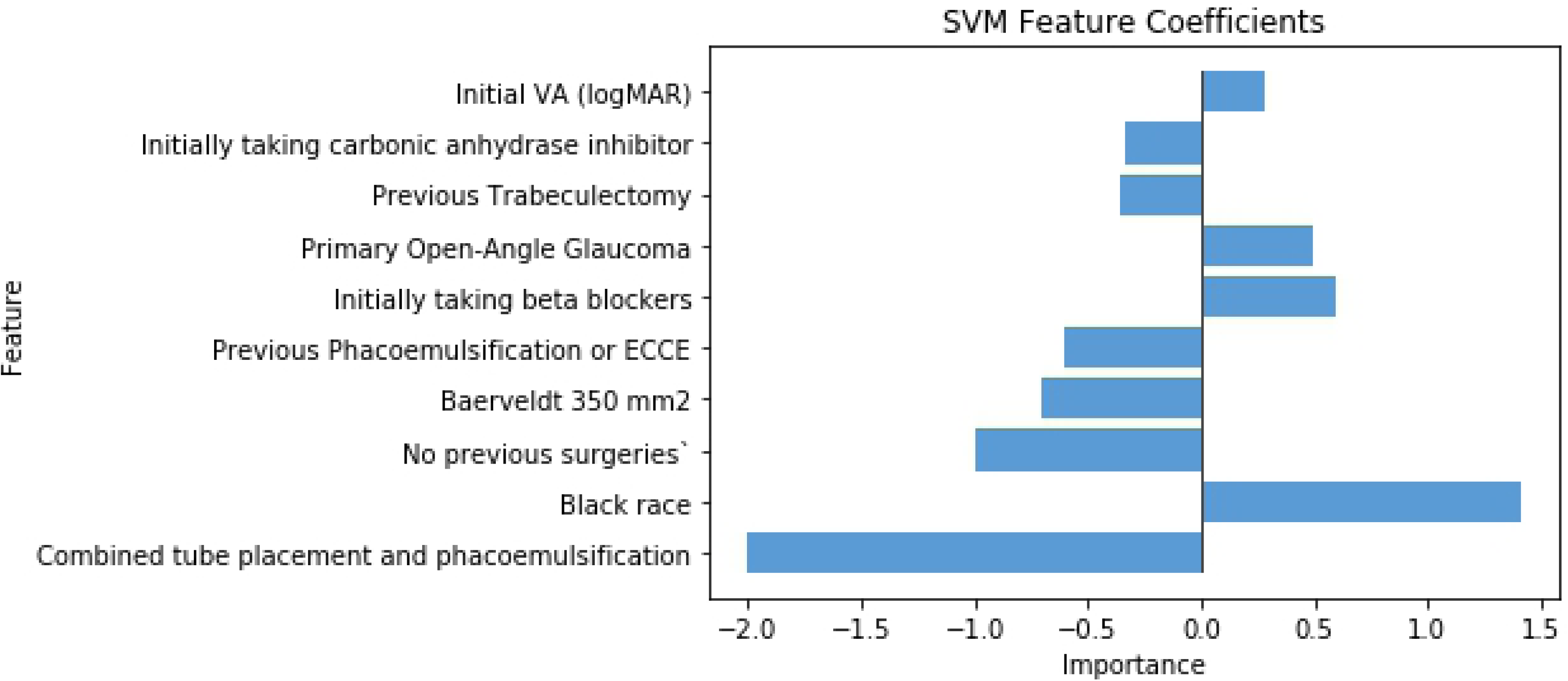
Feature Coefficients of SVM Classifier. The most significant factor promoting failure was black race, while the most significant factor mitigating failure was phacoemulsification performed in the same procedure as drainage device implantation.

Like regression, race and initially taking beta-blockers have the most weight in causing implant failure. Primary Open-Angle Glaucoma show possibility of implant failure too. The SVM classifier had an accuracy score of 0.61%±0.03 and an AUC value of 0.62±0.03.

### 3.3. Decision Tree

A decision tree repeatedly picks a threshold to divide data until it places all data items in groups (mostly) of the same class. First, it finds the threshold for all features dividing data most cleanly. Then it chooses features producing the cleanest split and repeats the process separately for the data on each side of the split. The algorithm stops when the data divide into pure groups or when the number of points in each group is too small to divide further without overfitting [13]. We used a minimum of three data points per leaf node and the Gini impurity measure.

As described in Table 5 and Fig 3, IOP and race are the most important factors in device failure. Table 6 and Fig 4 also show that many predicted outcomes depend on these features. Combined GDD placement and phacoemulsification, age, and usage of the 245 mm^2^ Molteno GDD are other significant factors. Unlike in the cases of regression and SVM, initially taking beta-blockers does not appear in decision tree.

**Table 5.** Decision Tree Feature Importances.

| Feature | Importance |
| --- | --- |
| Initial IOP | 0.19 |
| Black | 0.17 |
| In-Surgery Phacoemulsification | 0.16 |
| Age at Surgery | 0.14 |
| Molteno 245 mm<sup>2</sup> | 0.14 |
| Baerveldt 350 mm<sup>2</sup> | 0.07 |
| Initial VA (logMAR) | 0.06 |
| Primary Open-Angle Glaucoma | 0.04 |
| Initial Number of Medications | 0.02 |
| Previous Ex-Press Shunt | 0.01 |

**Table 6.** Patient Groups Created Using Decision Tree.

|  | Group Characteristics |  |  |  |  | Patient Features |  |  |  |  |  |  |  |  |  |
| --- | --- | --- | --- | --- | --- | --- | --- | --- | --- | --- | --- | --- | --- | --- | --- |
| Group Number | Decision | Count | Number of fairsuccesses | Number of failures | Gini | In-Surgery Phacoemulsification | Black | Age at Surgery | Molteno 245 mm² | Initial IOP | Primary Open-Angle Glaucoma | Previous Ex-Press Shunt | Initial Number of Medications | Initial VAllogMAR | Baerveldt 350 mm² |
| 0 | Success | 8 | 8 | 0 | 0 | False | Any | ≤ 75.5 | Any | Any | Any | Any | Any | Any | Any |
| 1 | Success | 3 | 2 | 1 | 0.4 | False | Any | > 75.5 | Any | Any | Any | Any | Any | Any | Any |
| 2 | Failure | 7 | 0 | 7 | 0 | True | False | Any | Any | Any | Any | Any | Any | Any | False |
| 3 | Success | 4 | 2 | 2 | 0.5 | True | False | Any | Any | Any | Any | Any | Any | Any | True |
| 4 | Failure | 4 | 0 | 4 | 0 | True | True | Any | False | Any | Any | Any | Any | ≤ 0.44 | Any |
| 5 | Success | 4 | 2 | 2 | 0.5 | True | True | Any | False | Any | Any | Any | Any | > 0.44 | Any |
| 6 | Failure | 3 | 0 | 3 | 0 | True | True | ≤ 73.5 | True | ≤ 15.75 | Any | Any | Any | Any | Any |
| 7 | Success | 4 | 3 | 1 | 0.4 | True | True | ≤ 73.5 | True | 21.0 ≤ x < 15.75 | False | Any | Any | Any | Any |
| 8 | Failure | 4 | 1 | 3 | 0.4 | True | True | ≤ 73.5 | True | 38.0 ≤ x < 21.0 | False | Any | Any | Any | Any |
| 9 | Success | 3 | 2 | 1 | 0.4 | True | True | ≤ 73.5 | True | > 38.0 | False | Any | Any | Any | Any |
| 10 | Success | 3 | 2 | 1 | 0.4 | True | True | ≤ 73.5 | True | > 15.75 | True | False | Any | Any | Any |
| 11 | Success | 4 | 4 | 0 | 0 | True | True | ≤ 73.5 | True | > 15.75 | True | True | ≤ 3.5 | Any | Any |
| 12 | Success | 3 | 2 | 1 | 0.4 | True | True | ≤ 73.5 | True | > 15.75 | True | True | > 3.5 | Any | Any |
| 13 | Success | 8 | 8 | 0 | 0 | True | True | > 73.5 | True | Any | Any | Any | Any | Any | Any |

**Fig 3.**
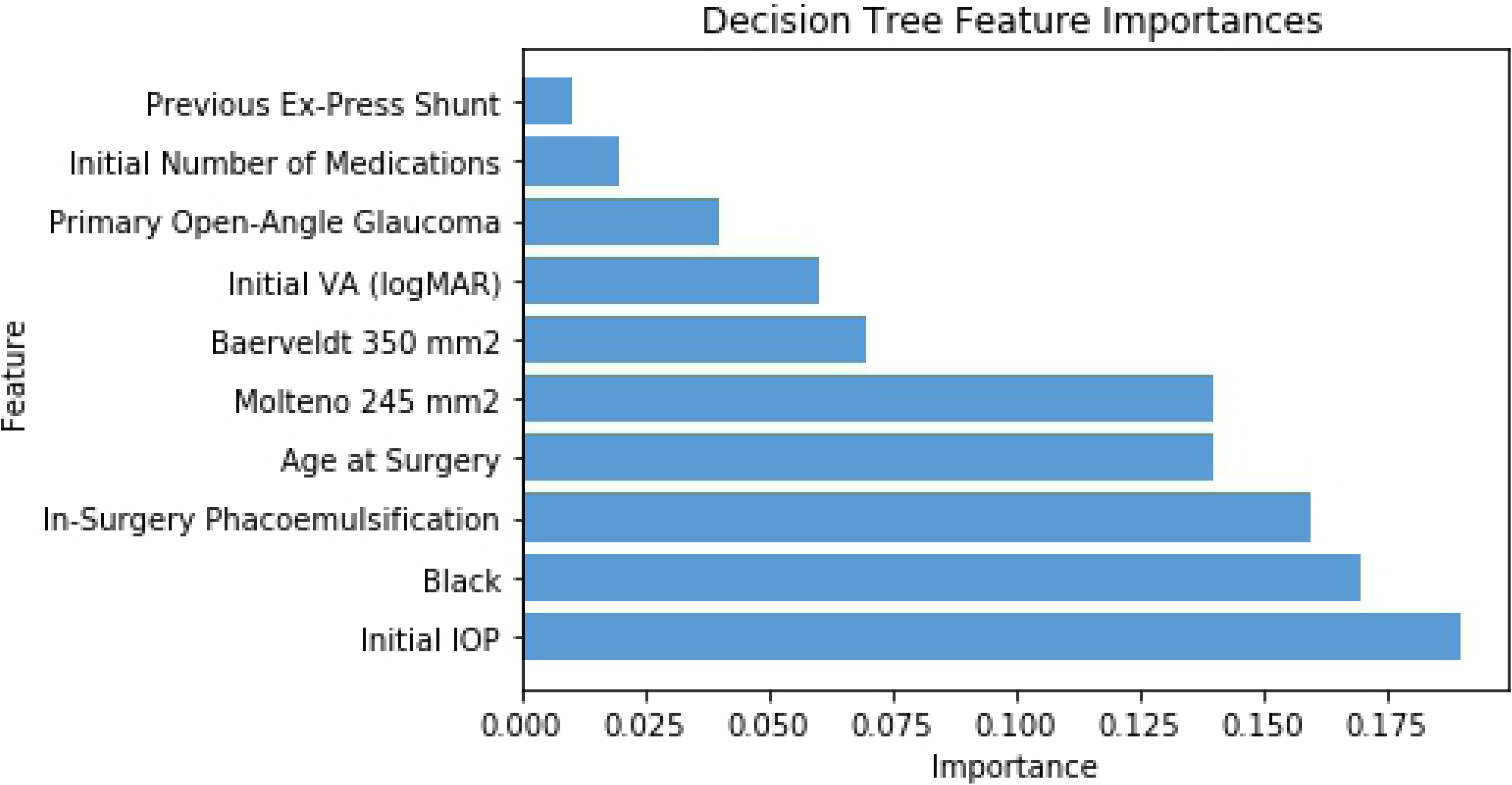
Decision Tree Feature Importances. The three most significant factors contributing to implant failure were the patient’s initial IOP, whether the patient was black, and whether phacoemulsification was performed in the same procedure as drainage device implantation.

**Fig 4.**
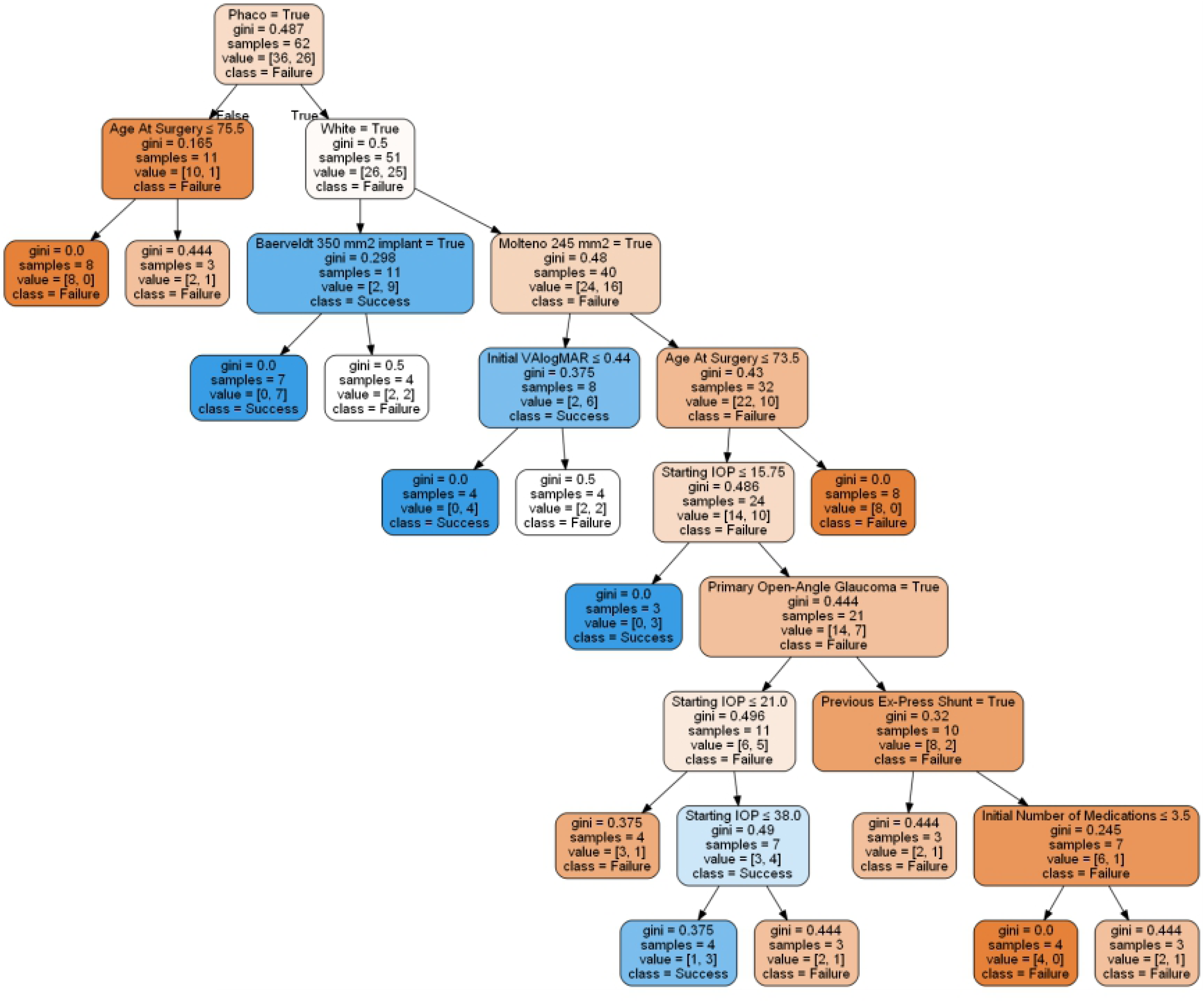
Decision Tree of Glaucoma Data. In deciding a patient’s outcome, the algorithm chooses which branch to follow based on the condition noted in the box.

The decision tree’s accuracy score was 0.5±0.05, and its AUC value was 0.45%±0.04. Low accuracy makes the efficiency of this method less than previous methods.

### 3.4. Artificial Neural Network

A feed-forward artificial neural network imitates biological neural tissue using sequential layers of “neurons” that transform the underlying data and pass it on to the next layer. The root of a neural net is a perceptron: two or more inputs connected to a neuron, which then multiplies each input by a weight, adds an intercept, and applies an output function to the result. In a single-layer neural network, many perceptrons extract information from the underlying features, and a final neuron (or more for multiclass classification) combines the output from these nodes [13]. A single-layer network with 25 hidden nodes (Fig 5) was trained on the data. The importance of each feature to this classifier is shown in Fig 6 and Table 7. Combined GDD placement and phacoemulsification was indicated as the most important factor in failure. Chronic angle-closure glaucoma and initially taking beta-blockers were also associated with therapy failure, though to a lesser extent. Race with importance = 4.6 shows high effect on failure. The accuracy score was 0.52±0.10 and the AUC value was .53±0.11.

**Table 7.** Importance of Features in Neural Network Classifier.

| Feature | Importance | Feature | Importance |
| --- | --- | --- | --- |
| In-surgery Phacoemulsification | 7.41 | No previous surgeries | 3.99 |
| Chronic angle-closure glaucoma | 5.81 | Open-Angle Glaucoma | 3.90 |
| Initially taking Beta Blockers | 5.17 | Previous Phacoemulsification or ECCE | 3.89 |
| Previous Express Shunt | 5.01 | Molteno 185 mm<sup>2</sup> | 3.87 |
| Baerveldt 350 mm<sup>2</sup> | 4.65 | Number of Previous Surgeries | 3.78 |
| Black | 4.56 | Age at Surgery | 3.54 |
| Combined Mechanism Glaucoma | 4.38 | Starting VAllogMAR | 3.53 |
| Molteno 245 mm<sup>2</sup> | 4.38 | Initially taking CAI | 3.30 |
| Initially taking Prostaglandins | 4.18 | Previous Trabeculectomy | 3.23 |
| Starting Number of Medications | 4.16 | White | 2.99 |
| Initially taking Alpha Agonists | 4.11 | Baerveldt 250 mm<sup>2</sup> | 2.99 |
| Male | 4.10 | Initial IOP | 2.98 |

**Fig 5.**
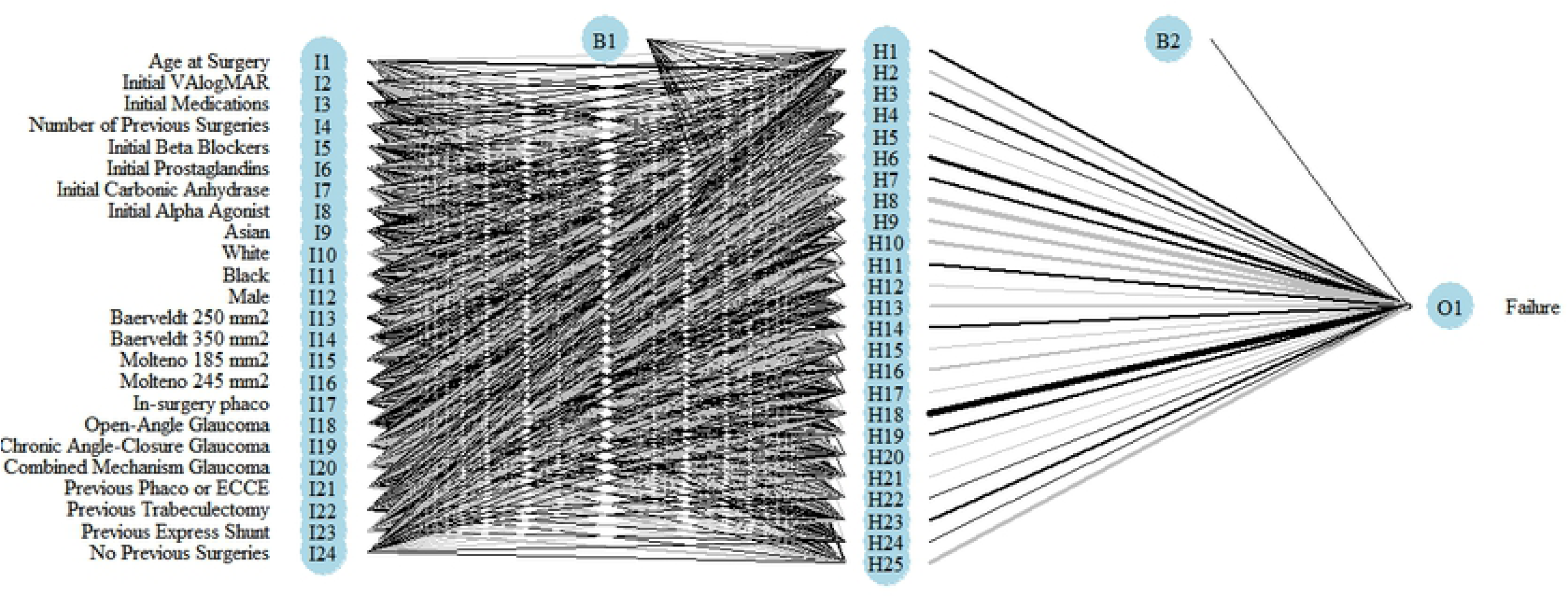
Neural Network Architecture.

**Fig 6.**
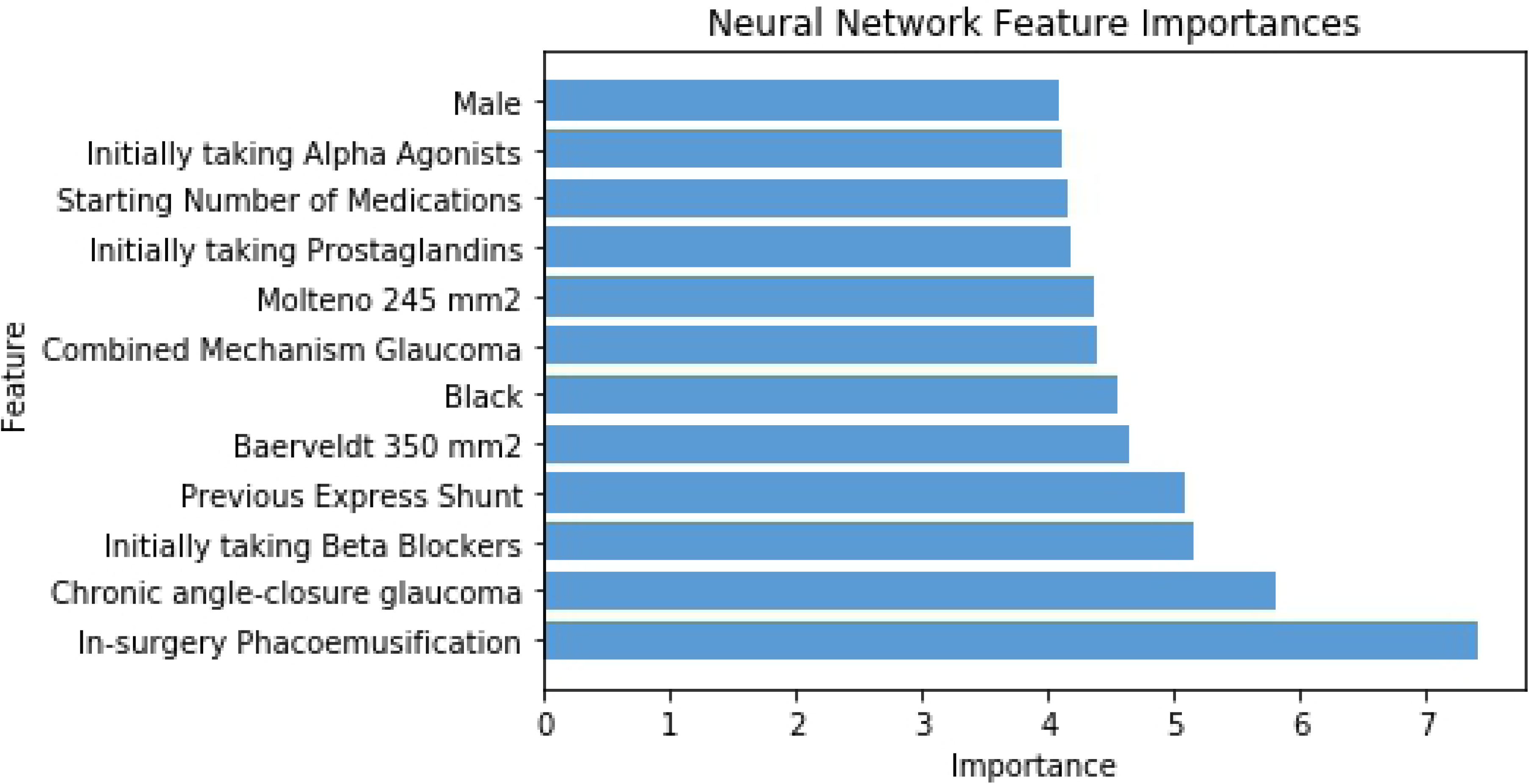
Neural Network Feature Importance. The three most important factors were whether phacoemulsification was performed in the same procedure, whether the patient had chronic angle-closure glaucoma, and whether the patient was initially taking beta blockers.

### 3.5. Random Forest

A random forest averages the predictions of multiple decision trees trained on subsets of the data [13]. Using ten decision trees, the algorithm identified age at surgery, initial IOP, and visual acuity as the most important factors determining device failure. Race and utilization of the larger Molteno device (245 mm^2^) were associated with device failure, though to a lesser degree. The overall AUC value was 0.58±0.1, and the accuracy score was 0.58±0.13. The importance of each feature to this classifier is shown in Table 8 and Fig 7.

**Table 8.** Importance of Features in Random Forest Classifier.

| Feature | Importance |
| --- | --- |
| Age at Surgery | 0.25 |
| Initial IOP | 0.2 |
| Initial VAllogMAR | 0.12 |
| Black | 0.07 |
| Molteno 245 mm<sup>2</sup> | 0.07 |
| Initial Number of Medications | 0.06 |
| On beta-blocker prior to surgery | 0.06 |
| Combined device and phacoemulsification | 0.06 |
| Number of Previous Surgeries | 0.05 |
| Primary Open-Angle Glaucoma | 0.03 |
| On CAI prior to surgery | 0.02 |
| Previous Ex-Press Shunt | 0.01 |

**Table 9.** Most Important Features across all models. Red: high effect, Orange: low effect

| Methods | Black race | Beta Blockers | Age | Initial IOP | Molteno<br>245 mm<sup>2</sup> | Cataract<br>removal |
| --- | --- | --- | --- | --- | --- | --- |
| Logistic Regression | Red | Red |  |  |  |  |
| SVM | Red | Red |  |  |  |  |
| Random Forest | Orange |  | Red | Red | Orange |  |
| Neural Network | Orange | Red |  |  | Orange | Red |
| Decision Tree | Red |  | Red | Red | Red | Red |

**Table 10.** Accuracy and ROC score across all models.

| **Methods** | **ROC AUC**<br>**(mean ± SD)** | **Accuracy**<br>**(mean ± SD)** |
| --- | --- | --- |
| **Logistic Regression** | 0.67±0.08 | 0.66±0.08 |
| **SVM** | 0.62±0.03 | 0.61±0.03 |
| **Random Forest** | 0.53±0.10 | 0.58±0.13 |
| **Neural Network** | 0.52±0.10 | 0.53±0.11 |
| **Decision Tree** | 0.45±0.04 | 0.50±0.05 |

**Fig 7.**
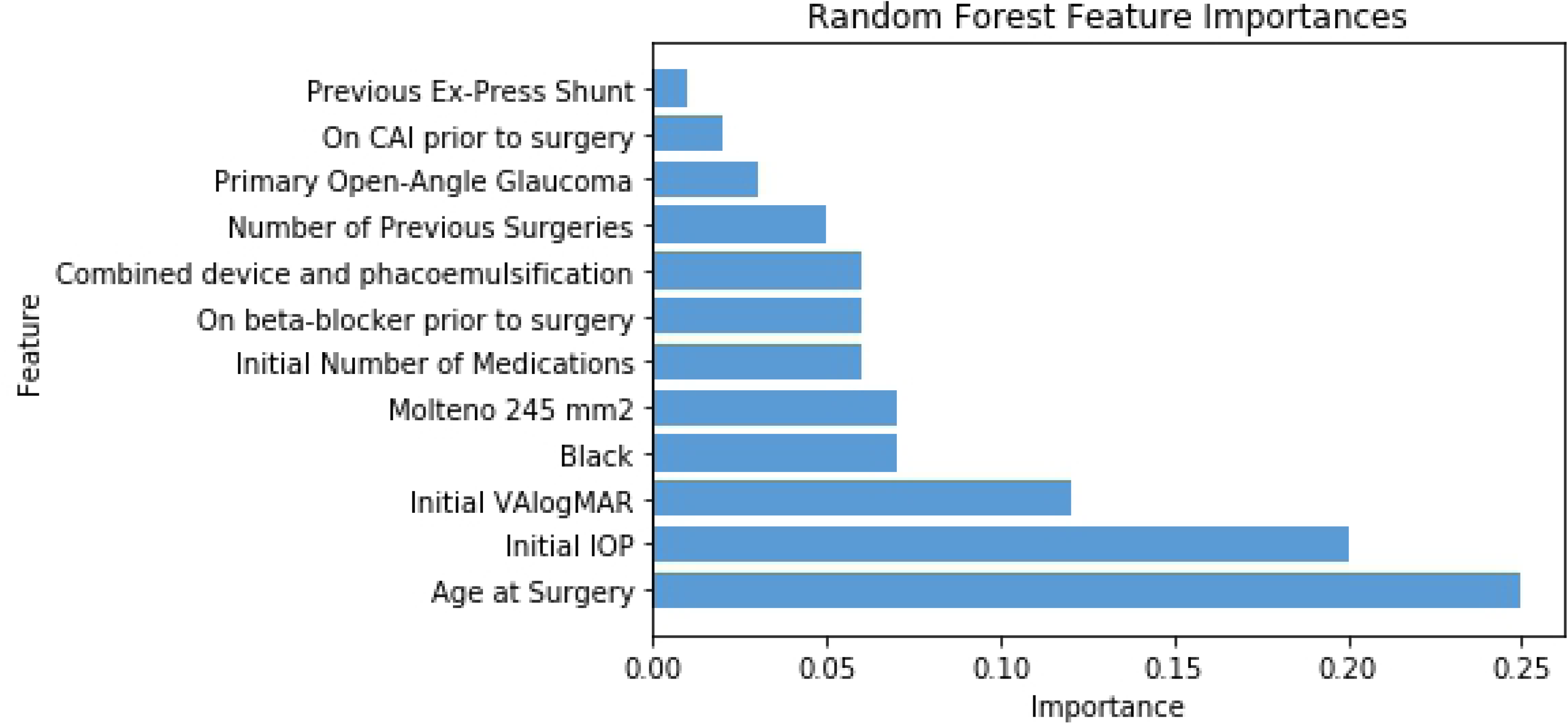
Feature Importance of Random Forest Classifier. The three most important factors were age at surgery, initial intra-ocular pressure, and initial visual acuity.

## 4. Conclusion

Comparing results from different models identified Black race as the strongest factor associated with device failure. This finding aligns with existing research in the ophthalmology literature [14]. Such failure rates are believed to be due to genetic differences in wound healing and proliferation of fibrovascular tissue [15]. Use of topical beta-blockers pre-operatively was also associated with device failure. In treating glaucoma medically, prostaglandin analogs are often first-line with beta-blockers used second-line or as a reasonable alternative first-line agent. Alpha-agonists and carbonic anhydrase inhibitors are often added next, though they can cause intolerable allergic reactions and discomfort on instillation, respectively [16]. These side effects can lead to drop intolerance and serve as an impetus for surgery. Therefore, it is perhaps not unsurprising that patients would be on beta-blockers when surgical intervention is needed as they are usually well-tolerated in those without respiratory problems. Nonetheless, beta-blockers association with implant failure in several models may be an area of further investigation. Placement of the larger Molteno GDD was associated with device failure, though this was a weaker association and found in weaker models. Again, this warrants further investigation given the devices function similarly. Lastly, age, increased IOP, and phacoemulsification at the time of GDD implantation were associated with failure in weaker models. Overall, the most accurate model was logistic regression, followed by a support vector machine model with a linear kernel. Our findings suggest machine learning techniques can accurately determine important features leading to failure of GDD implants from a large dataset of common clinical descriptors.

